# RNA virus discovery in Australian camelids reveals divergent picornaviruses and the convergent evolution of upstream ORFs

**DOI:** 10.64898/2026.02.19.706906

**Authors:** Kosuke Takada, Jonathon C. O. Mifsud, Junki Hirano, Erin Harvey, Sabrina Sadiq, Bethan J. Lang, Yoshiharu Matsuura, Edward C. Holmes

## Abstract

Invasive species can impact viral ecology, evolution and emergence by acquiring and disseminating viruses absent from native hosts. However, the extent to which invasive species harbour previously unrecognized RNA viruses and transmit these to native species is uncertain. We performed metatranscriptomic sequencing of invasive camelids in Australia and identified several previously undescribed vertebrate-associated RNA viruses, including an astrovirus closely related to avian-associated viruses suggesting a recent host jump. We also identified highly divergent picornaviruses that differed sufficiently from recognized taxa in genome organization and polyprotein phylogeny to establish a new genus. Notably, one virus encoded a putative upstream ORF (uORF) in the 5′ genomic region. Across the *Picornaviridae*, putative uORF gain and loss appear to have occurred multiple times independently. In addition, although most of these uORF-encoded proteins exhibited little to no amino acid sequence homology, a subset showed overlapping ranges of secondary structure composition and intrinsic disorder, and when heterologously expressed, these proteins were translated and triggered reproducible transcriptional responses in a cell line. While no single pathway was uniformly affected across all uORFs, distinct uORFs from divergent lineages consistently perturbed overlapping sets of cellular pathways, supporting broadly analogous functional effects despite a lack of sequence homology. These findings demonstrate that uORFs represent a recurrent and selectable functional module within RNA virus genomes, suggest that the upstream genomic position itself constitutes a “hotspot” for the repeated acquisition of a functional module, and provide experimental evidence that their functional properties have converged across evolutionarily independent lineages.

**Author Summary:** RNA viruses evolve under strong genomic constraints, forcing them to repeatedly adopt similar solutions to common challenges posed by their host environment. Invasive species can impact viral ecology, evolution and emergence by acquiring and disseminating non-native viruses. By characterizing RNA viruses infecting invasive camelids in Australia, we discovered previously unrecognized RNA viruses and recurrent patterns of genome evolution. Notably, we identified a small additional gene located at the 5′ genomic region, known as an upstream open reading frame (uORF). uORFs were independently acquired and lost across multiple viral lineages, revealing that the same genomic region can be repeatedly exploited to acquire new functions. Although uORF-encoded proteins share little or no amino acid sequence homology, some of the encoded proteins showed comparable secondary structure composition and induce overlapping host cellular responses when expressed in cells. Hence, RNA viruses that have followed different evolutionary paths can converge on similar functional strategies.

## Introduction

The geographic isolation of the Australian continent, which separated from eastern Antarctica approximately 38–45 million years ago, has led to the evolution of unique animals and plants [1]. Indeed, Australia is known for its biodiversity and abundance of endemic species [1]. In such an isolated environment, the introduction and impact of invasive species has been particularly evident, and animals introduced since the 19th century have had major injurious economic and ecological consequences in Australia, with the best documented example being the destructive continental spread of the European rabbit (*Oryctolagus cuniculus*) [2]. Invasive species not only compete with native species for resources and threaten biodiversity, but also serve as reservoirs and transmission conduits for viruses and other pathogens [3, 4]. Hence, the introduction of non-native species has the potential to facilitate the emergence or spread of infectious diseases, further contributing to the disruption and collapse of existing ecosystems.

Invasive species are also often characterised by major demographic changes, particularly population bottlenecks, following introduction that can alter viral population structure and facilitate the emergence of genetically divergent lineages. For instance, dromedary camels (*Camelus dromedarius*) were introduced from India and Afghanistan in the 18th century for the inland exploration and transport of goods, and have since become feral [5], with a current minimum population estimate of approximately 1 million feral camels now living predominantly in Australia [6]. Up to 20,000 dromedary camels were introduced between 1880 and 1907 and their population grew rapidly, exceeding 300,000 by the early 2000s [6]. To address the resulting environmental and cultural damage, a large-scale culling program between 2008 and 2013 reportedly reduced their numbers to around 300,000, although this figure is disputed [7]. Alpacas (*Vicugna pacos*) and llamas (*Lama glama*) were first introduced into Australia from South America in the 19th century, but did not become established at that time [8]. They were more successfully introduced during the 20th century, and are now bred primarily for ornamental purposes and for their high-quality fibre [9, 10]. Together, these historic introductions have generated host populations shaped by repeated bottlenecks, founder effects, and ecological perturbation, creating conditions that may accelerate viral emergence. However, the emergence of novel viral lineages in invasive host populations also provides an opportunity to investigate how genomic innovations arise and persist under changing ecological and demographic conditions.

Recent studies have shown that RNA viruses from multiple families (i.e., the *Coronaviridae*, *Filoviridae*, *Flaviviridae, Picornaviridae* and *Retroviridae*) encode upstream open reading frames (uORFs) that influence viral replication and tissue tropism [11–16]. However, these elements often lack detectable sequence conservation and exhibit patchy distributions among related taxa, obscuring their origins and functional significance. Notably, because many RNA virus genomes recovered from metagenomic studies are incomplete, particularly at the 5′ termini where uORFs occur, their apparent patchy distribution may in part reflect detection biases rather than true evolutionary loss. Consequently, it is unclear whether uORFs arise sporadically through lineage-specific innovation or represent recurrent, selectable modules that are repeatedly gained and lost during viral evolution. Resolving this question requires the discovery and comparative analysis of diverse viral lineages, for example in ecological contexts that may foster rapid diversification such as species invasions. Consequently, identifying novel RNA viruses from ecologically perturbed systems may provide critical opportunities to investigate how uORFs emerge, diversify, and are maintained.

Despite major advances in virus metagenomics, the extent to which non-native species harbour previously unrecognized RNA viruses remains uncertain. Herein, we describe a metatranscriptomic analysis of invasive camelids (alpacas, camels and llamas) in Australia to characterize their RNA viromes and identify novel virus lineages that in turn provide insights into RNA virus genome evolution.

## Results

### Sample collection and virome analysis of non-native Australian camelids

We determined the virome of camels, llamas, and alpacas sampled from locations across Australia (Fig 1A). Total RNA sequencing was performed on 154 alpaca samples (pooled into 25 libraries), 118 camel samples (pooled into 26 libraries) and 14 llama samples (pooled into 5 libraries), with pooling based on geographic location and sample type (Fig 1B and Supplementary Table S1). *De novo* assembly of the sequencing reads yielded a median of 327,207 contigs per library for alpacas (range: 195,425-677,502), 281,879 for camels (range: 265-659,24) and 441,517 for llamas (range: 264,769-592,016), resulting in a total of 5,056,108, 2,220,017, and 9,433,381 contigs, respectively (Supplementary Table S1).

**Fig 1.**
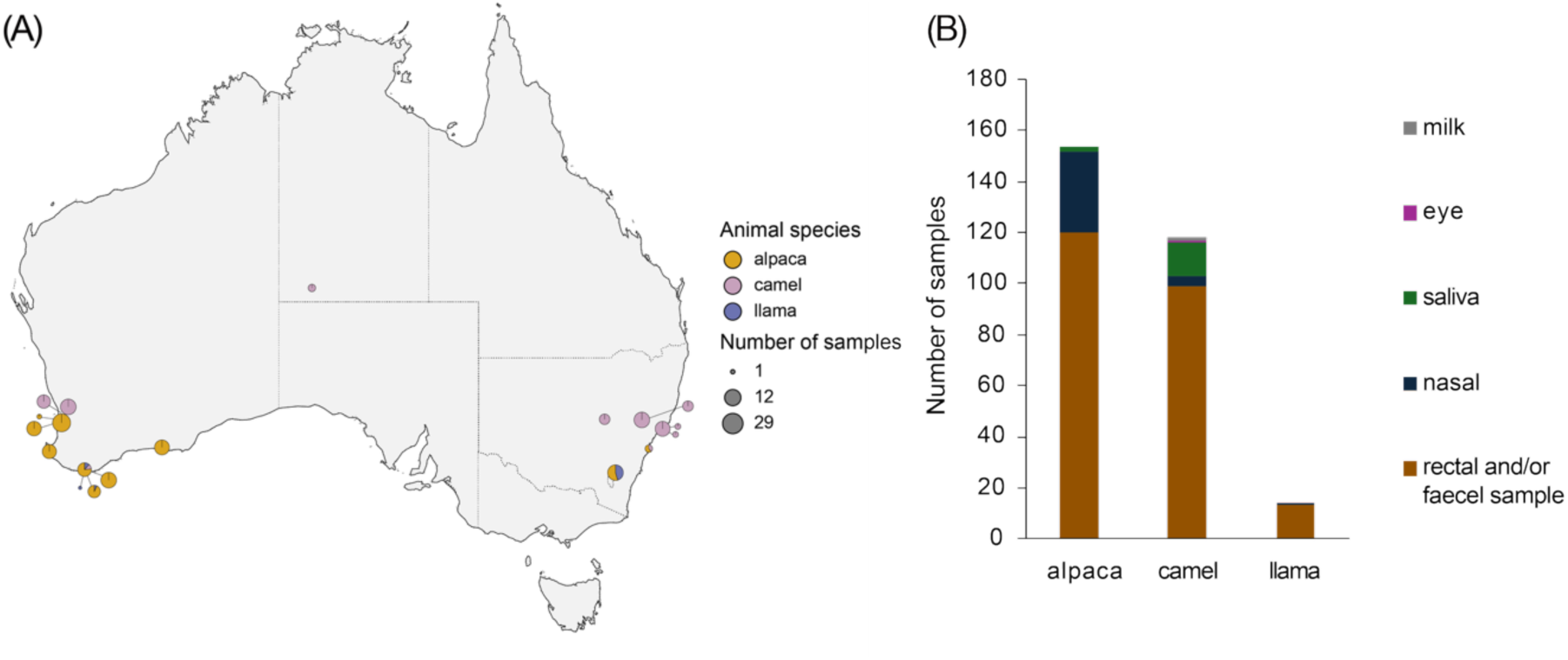
Geographic location and sample types of alpaca, camel and llama-derived samples. (A) Map showing the composition and number of the alpaca, camel and llama samples across mainland Australia. The circles and pie chart colours reflect animal species, and their size reflects the number of individual samples collected at each location; representative sizes are shown in the legend. (B) Sample types collected from each animal species.

We focused on RNA viruses known to infect vertebrates, excluding those likely associated with diet or gut microbiota such as picobirnaviruses [17]. In total, we identified five vertebrate-associated RNA viruses in alpaca and camel faecal samples (Table 1). No RNA viruses were identified in the llama samples. These viruses were classified into four species, three of which represent novel species within the family *Picornaviridae*. The remaining virus belonged to the *Astroviridae*. Consistent with findings from other viral metagenomic studies in mammals, nearly all the viruses identified in this study were novel, with the exception of one picornavirus that showed close similarity to a previously reported camel-associated virus from Eurasia. Notably, no coronaviruses, including Middle East respiratory syndrome–related coronaviruses (MERS-like CoVs), were identified in any of the Australian camelid samples.

**Table 1.**
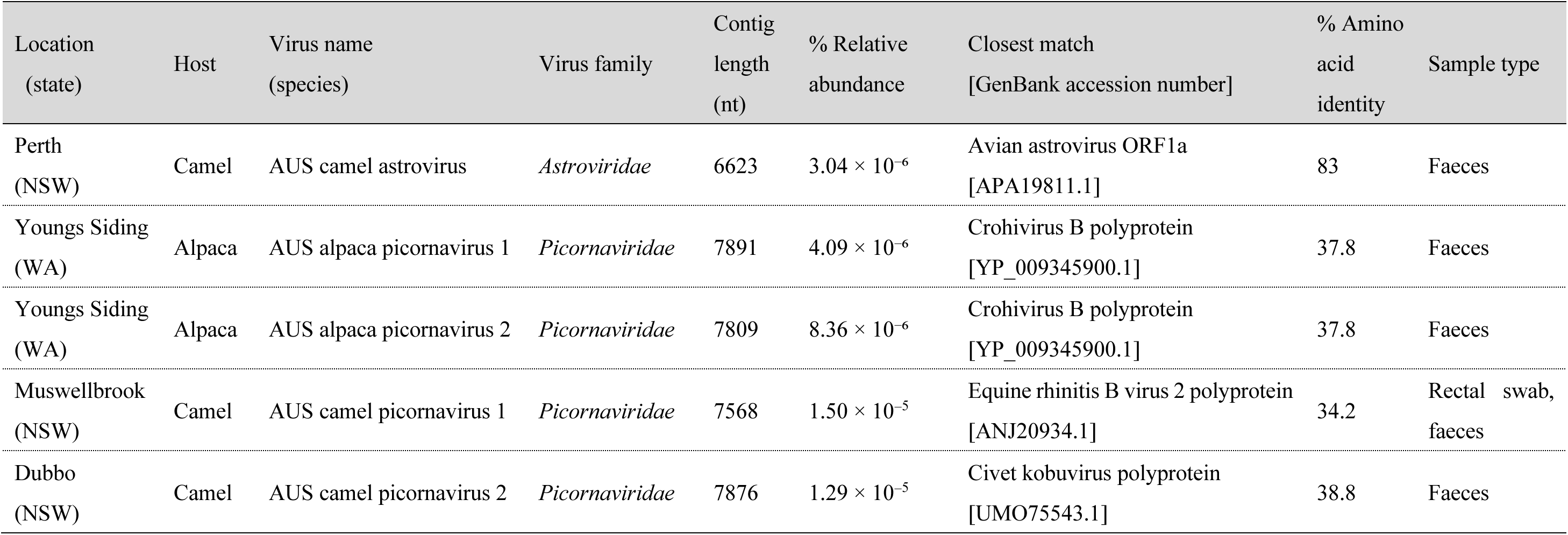
Vertebrate-associated RNA virus contigs, contig length (nt), percentage abundance in their respective pools, and percentage amino acid identity to their closest match on NCBI/GenBank.

| Location<br>(state) | Host | Virus name<br>(species) | Virus family | Contig<br>length<br>(nt) | % Relative<br>abundance | Closest match<br>[GenBank accession number] | % Amino<br>acid<br>identity | Sample type |
| --- | --- | --- | --- | --- | --- | --- | --- | --- |
| Perth<br>(NSW) | Camel | AUS camel astrovirus | *Astroviridae* | 6623 | $3.04 \times 10^{-6}$ | Avian astrovirus ORF1a<br>[APA19811.1] | 83 | Faeces |
| Youngs Siding<br>(WA) | Alpaca | AUS alpaca picornavirus 1 | *Picornaviridae* | 7891 | $4.09 \times 10^{-6}$ | Crohivirus B polyprotein<br>[YP_009345900.1] | 37.8 | Faeces |
| Youngs Siding<br>(WA) | Alpaca | AUS alpaca picornavirus 2 | *Picornaviridae* | 7809 | $8.36 \times 10^{-6}$ | Crohivirus B polyprotein<br>[YP_009345900.1] | 37.8 | Faeces |
| Muswellbrook<br>(NSW) | Camel | AUS camel picornavirus 1 | *Picornaviridae* | 7568 | $1.50 \times 10^{-5}$ | Equine rhinitis B virus 2 polyprotein<br>[ANJ20934.1] | 34.2 | Rectal swab,<br>faeces |
| Dubbo<br>(NSW) | Camel | AUS camel picornavirus 2 | *Picornaviridae* | 7876 | $1.29 \times 10^{-5}$ | Civet kobuvirus polyprotein<br>[UMO75543.1] | 38.8 | Faeces |

### Vertebrate-associated RNA viruses detected in camel and alpaca faecal samples

#### Astroviridae

We detected a novel astrovirus (positive-sense single-stranded RNA virus), provisionally named AUS camel astrovirus, in one of the pooled libraries generated from multiple camel faecal samples. Based on its ORF structure, the genome appears to be putatively near-complete (Fig 2A). Notably, the sequence shared 83% amino acid identity with Avian astrovirus isolate D11-440/HK ORF1a (GenBank accession APA19811.1) and had a relative abundance of 3.04 × 10⁻⁶ %, calculated as the percentage of total library reads (Table 1). Phylogenetic analysis of the RdRp and capsid proteins revealed that this virus grouped with chicken astrovirus within a broader group of avian-associated astroviruses and not with the mammalian astroviruses as might be expected (Fig 2B and 2C). Interestingly, a similar phylogenetic pattern was observed for the mammalian sequences Avastrovirus sp. isolate astro_vt_80 [(PP871798.1) from Viet Nam (wild boar, *Sus scrofa*) and Fox astrovirus isolate Pl1.4 (PV999258.1) from Italy (red fox, *Vulpes vulpes*), both of which also grouped within the avian-associated astrovirus clade based on analyses of the RdRp and capsid proteins, although in different phylogenetic positions to the camel sequences (Fig 2B-E). More broadly, it was notable that sequences of the avian genus *Avastrovirus* fell within the phylogenetic diversity of mammalian viruses.

**Fig 2.**
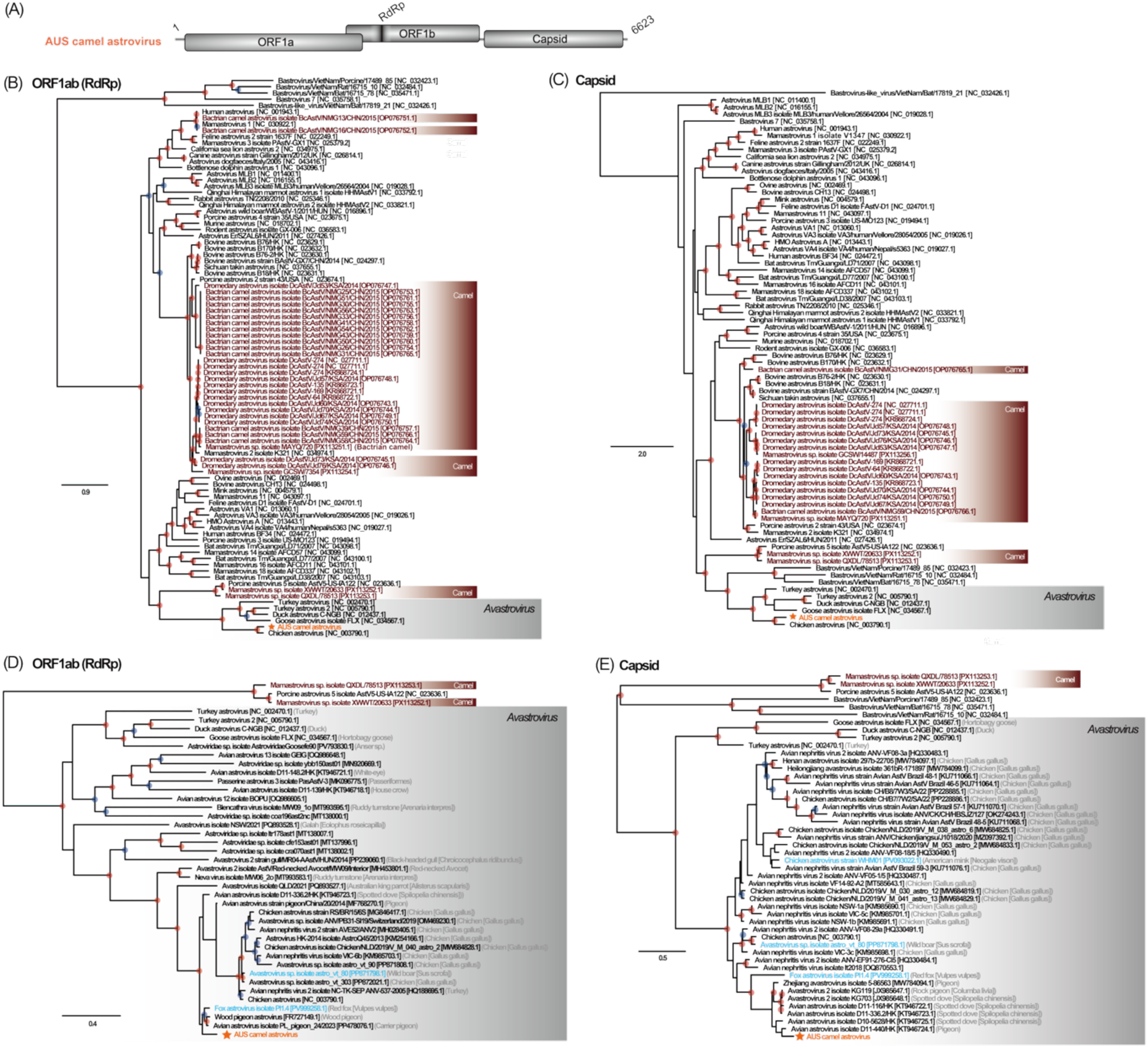
Phylogenetic trees of a novel virus belonging to the *Astroviridae*. (A) Schematic overview of the newly identified camel astrovirus. Predicted protein domains identified in the novel viral genomes using BLASTX are shown. Details are provided in Supplementary Table S2. (B–C) Maximum likelihood phylogenetic trees inferred from amino acid sequences of ORF1ab (RNA-dependent RNA polymerase; B) and ORF2 (capsid protein; C), including viruses identified in this study and representative reference viruses. Sequences classified as members of the genus *Avastrovirus* (i.e. from birds) are highlighted in grey. Sequences derived from camels outside Australia are highlighted in brown. The light blue labels indicate viruses detected from mammals that cluster within the *Avastrovirus* lineage. (D-E) Maximum likelihood phylogenies inferred from realigned amino acid alignments of avastrovirus and the related camel viruses. Branch lengths indicate the number of substitutions per site. Red and blue circles at internal nodes represent bootstrap values ≥ 90% and ≥ 80%, respectively. The global phylogenetic tree was mid-point rooted. For the magnified views focusing on specific clades, including sequences within the *Avastrovirus* lineage, *Mamastrovirus* sequences were used as an outgroup to provide directional context.

To determine whether the astrovirus identified here was truly associated with camels, we used Kraken to analyse the distribution of reads in the library. As expected from a faecal metatranscriptomic data set, most reads were derived from bacteria (52.7%) and archaea (29.6%), with eukaryotic reads representing a minor proportion (17.1%) of the data. However, within the eukaryotic fraction, reads assigned to Mammalia accounted for 1.14% of the total data set, including 0.64% assigned to Artiodactyla (mammals), whereas avian reads represented only 0.07% (Supplementary Fig S1). Hence, this virus was likely associated with camel hosts.

#### Picornaviridae

Several picornaviruses (positive-sense single-stranded RNA viruses) were also identified in camels and alpacas. Two crohivirus-related sequences, putatively near-complete based on ORF structure, were detected in the alpaca faecal samples (Fig 3A). The two genomes identified here were identical except for minor differences in terminal lengths, resulting in an identical polyprotein amino acid sequence and tight clustering in the phylogeny (Fig 3B and 3C). The longer sequence, provisionally named AUS alpaca picornavirus 1, had highest amino acid identity with Crohivirus B clone Bat/CAM/CroV-P25/2013 polyprotein [YP_009345900.1] (37.8%) detected in *Eidolon helvum* feces from Cameroon, and had a relative abundance of 4.09 × 10⁻⁶ % (Table 1). However, phylogenetic analysis of the polyprotein showed that the sequence was most closely related to that from a virus detected through the metagenomic analysis of the enteric RNA virome of calves in Ethiopia (Fig 3C and Supplementary Fig S2), and the relatively high genetic divergence from other crohiviruses suggests that this sequence may represent distinct evolutionary lineage.

**Fig 3.**
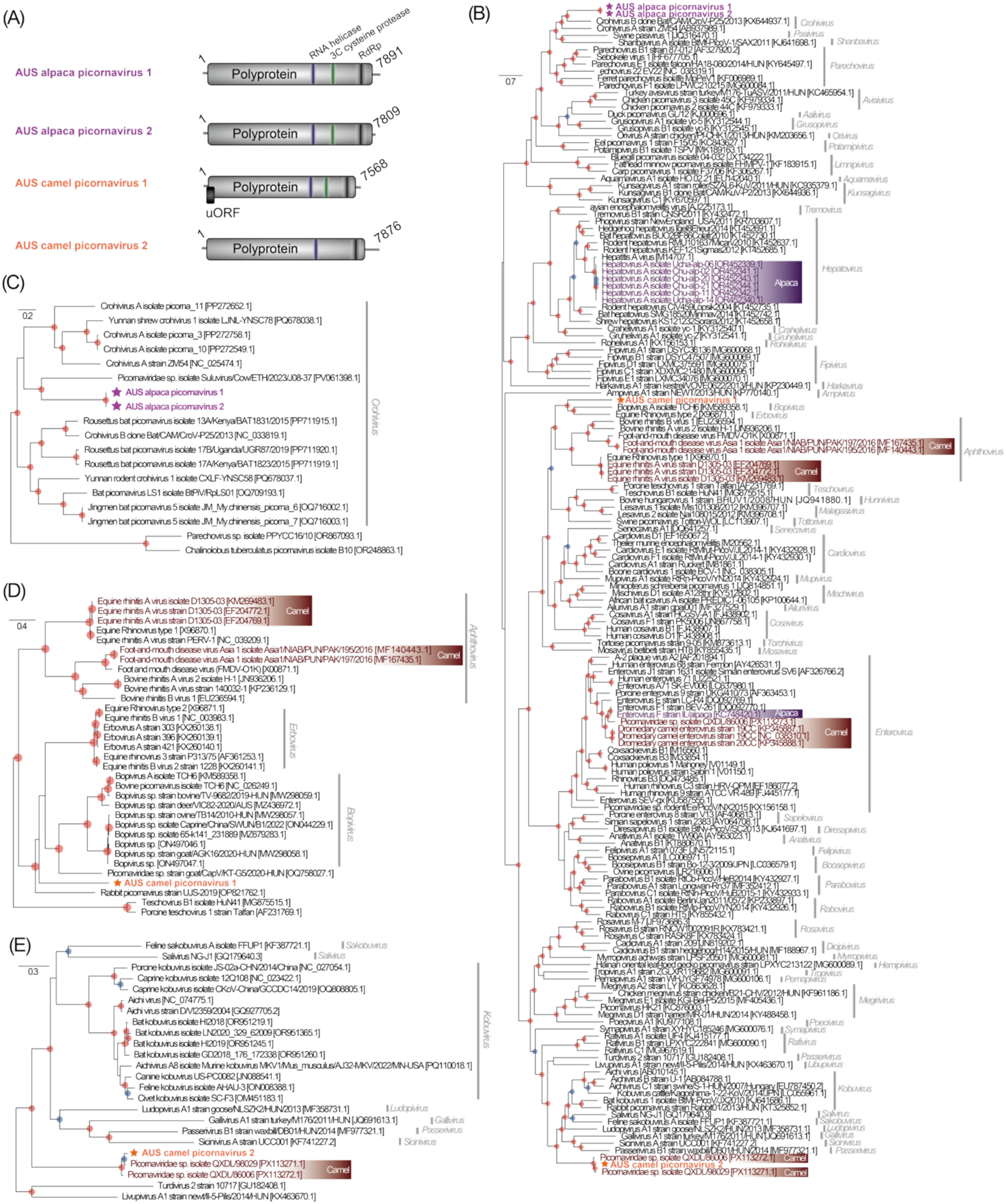
Phylogenetic trees of four viruses belonging from the *Picornaviridae*. (A) Schematic overview of the newly identified picornaviruses. Predicted protein domains identified in the novel viral genomes using BLASTX are shown. Details are provided in Supplementary Tables S3-S6. (B) Maximum likelihood phylogenetic tree inferred from amino acid sequences of the polyprotein, including viruses identified in this study and representative reference viruses. Sequences derived from camels outside Australia are highlighted in brown, and those derived from alpacas are highlighted in purple. (C–E) Maximum likelihood phylogenies inferred from realigned amino acid sequences of subsetted taxa corresponding to (C) *Crohivirus*, (D) *Erbovirus*/*Bopivirus*/*Aphthovirus*, and (E) *Kobuvirus*/the other genera. Branch lengths represent the number of substitutions per site. Red and blue circles at internal nodes indicate bootstrap values ≥ 90% and ≥ 80%, respectively. All phylogenetic trees were midpoint rooted for clarity only.

Another picornavirus sequence, putatively near-complete based on genome organization, was identified in a pooled library generated from multiple camel faecal samples (Fig 3A). This sequence, provisionally named AUS camel picornavirus 1, had 34.2% amino acid identity with the Equine rhinitis B virus 2 1228 polyprotein [ANJ20934.1] with a relative abundance of 1.50 × 10^⁻5^ % (Table 1). Based on analysis of the polyprotein, the virus grouped within a broader clade comprising rabbit picornavirus, erbovirus (i.e., equine rhinitis B virus) and bovine picornavirus (i.e., bopivirus) as well as goat picornaviruses (Fig 3B and 3D). However, it formed a distinct and well-supported lineage that was clearly separated from each of these viruses at the genus level, indicating that this virus likely represents a new picornavirus genus. Interestingly, this sequence was not closely related to picornaviruses detected from *Camelus dromedarius* in the United Arab Emirates (D1305-03 [KM269483.1], D1305-03 [EF204772.1], and D1305-03 [EF204769.1]) classified as Equine rhinitis A virus, and from *Camelus dromedarius* in Pakistan (Foot-and-mouth disease virus [FMDV] Asia 1 isolates Asia1/NIAB/PUN/PAK/195/2016 [MF140443.1] and Asia1/NIAB/PUN/PAK/197/2016 [MF167435.1]) from the genus *Aphthovirus*). Similarly, Bopivirus sp. deer/VIC82-2020/AUS [MZ436972.1], which was detected from deer in Australia, was phylogenetically distinct from the AUS camel picornavirus 1 sequence detected here.

Finally, a kobuvirus sequence was detected in one of the pooled libraries generated from multiple camel faecal samples (Fig 3A). This sequence, provisionally named AUS camel picornavirus 2, exhibited 38.8% amino acid identity with civet kobuvirus isolate SC-F3 polyprotein [UMO75543.1] (Table 4), and phylogenetic analysis of the polyprotein revealed that it was most closely related to other mammalian kobuviruses (Fig 3B and 3E). Notably, although only a partial sequence of the capsid protein and RdRp was detected here, it clustered with Picornaviridae sp. isolates QXDL/98029 [PX113271.1] and QXDL/86006 [PX113272.1], both of which were detected in Chinese Bactrian camels (Fig 3B and 3E and Supplementary Fig S3). Hence, this kobuvirus sequence likely represents a camel-specific lineage, and potentially a novel genus, that was present in the founding camel population introduced to Australia.

### A putative upstream open reading frame (uORF) predicted in the AUS camel picornavirus 1 genome

Since enterovirus genomes possess a polyprotein encoded in a single open reading frame and also harbor a uORF that is under strong purifying selection [13], the presence of a putative uORF in the AUS camel picornavirus 1 genome was noteworthy (Fig 3A and 4A). To more precisely document the phylogenetic position of the AUS camel picornavirus 1, we estimated phylogenetic trees for individual amino acid alignments of the 3CD protein (i.e., the precursor comprising 3C cysteine protease and RNA-dependent RNA polymerase: RdRp), 3D protein (i.e., RdRp) and P1 region (encoding L, VP4, VP2, VP3 and VP1), together with representative viruses closely related to AUS camel picornavirus 1 (i.e., *Erbovirus*, *Bopivirus*, and *Aphthovirus*). We performed the same analyses for the other novel picornaviruses identified in this study (Supplementary results). Phylogenetic analyses of the structural (i.e., P1 region) and non-structural (i.e, 3CD protein and 3D protein) proteins (Fig 4A and Supplementary Table S9) yielded consistent, but lineage-specific, patterns (Fig 4B–4D). Specifically, in the 3D tree, the AUS camel picornavirus 1 sequence clustered with the genus *Erbovirus* (Fig 4B). In marked contrast, in the 3CD phylogeny the AUS camel picornavirus 1 sequence fell within a clade containing both *Bopivirus* and *Erbovirus*, while in the P1 tree it fell within a broader clade comprising *Aphthovirus*, *Bopivirus*, and *Erbovirus* (Fig 4C and 4D). The 3D protein of AUS camel picornavirus 1 showed the highest amino acid sequence identity (43.38%) to Equine rhinitis B virus 1 [NC_003983.1], with the BLASTp hit covering over 85% of the query sequence. Taken together, these data confirm that AUS camel picornavirus 1 is sufficiently genetically distinct to represent a novel genus within the *Picornaviridae*.

**Fig 4.**
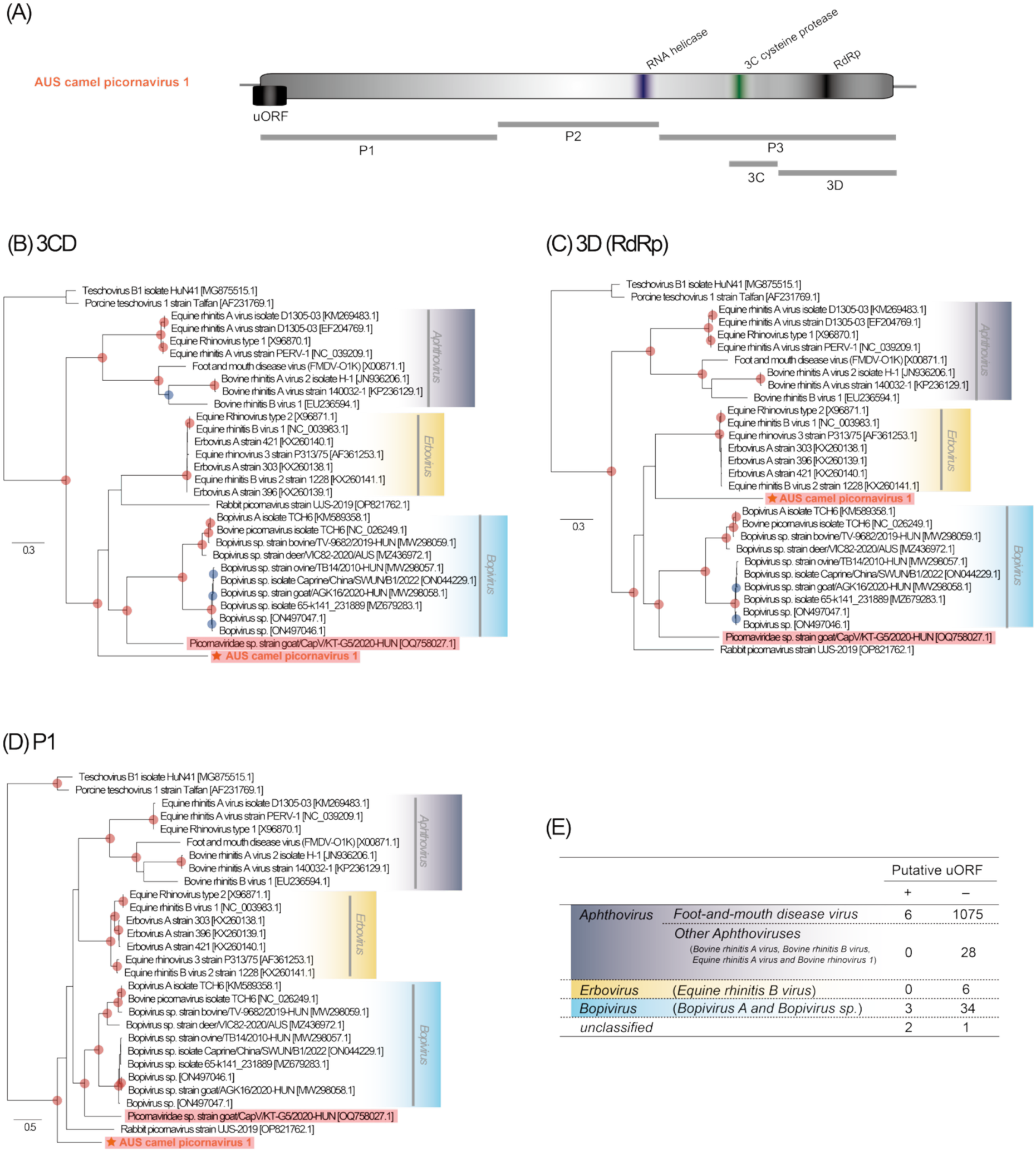
Individual gene phylogenies for the AUS camel picornavirus 1. (A) Schematic overview of structural (P1 region) and non-structural (2A–3D proteins) genes in AUS camel picornavirus 1. (B-D) Maximum likelihood phylogenetic trees based on amino acid sequences of the (B) 3CD and (C) 3D (RdRp) proteins, and (D) P1 region, including the AUS camel picornavirus 1 and representative *Erbovirus*, *Bopivirus*, and *Aphthovirus* sequences. Viruses with putative uORF are highlighted in red. Branch lengths represent the number of substitutions per site. Red and blue circles at internal nodes indicate bootstrap values ≥ 90% and ≥ 80%, respectively. Based on their phylogenetic positions inferred from the polyprotein tree (Fig 3), Teschovirus B1 isolate HuN41 (MG875515.1) and Porcine teschovirus 1 Talfan (AF231769.1) were used as outgroups to root the 3CD, 3D, and P1 phylogenetic trees. (E) Number of viruses with putative uORFs in each virus genus.

Next, we investigated those complete virus genome sequences that contain putative uORFs at similar genomic positions in virus genera closely related to AUS camel picornavirus 1 (i.e., *Erbovirus*, *Bopivirus*, and *Aphthovirus*). In the genus *Erbovirus*, none of the six sequences contained viruses with putative uORFs, whereas in *Bopivirus* and *Aphthovirus* three of the 37 sequences and six of the 1,124 sequences contained viruses with putative uORFs, respectively (Fig 4E and Supplementary Table S10). In addition, Picornaviridae sp. goat/CapV/KT-G5/2020-HUN (OQ758027.1), which is closely related to AUS camel picornavirus 1, has a putative uORF, while Rabbit picornavirus UJS-2019 (OP821762.1) does not (Fig 4B-E and Supplementary Table S10). Interestingly, no viruses with putative uORFs were found among respiratory-tropic viruses (i.e., Bovine rhinitis B virus, Equine rhinitis A virus and Equine rhinitis B virus), which predominantly infect the respiratory tract (Fig 4E and Supplementary Table S10). This is similar to previous findings showing that uORFs are rarely found in rhinoviruses, which are members of the genus *Enterovirus* and target the respiratory tract [13].

Finally, we investigated whether representative viruses from the broader *Picornaviridae* possess a putative uORF at a similar genomic position. This revealed that viruses carrying a putative uORF are distributed across phylogenetically disparate lineages, without forming a single monophyletic group (Fig 5A). In addition, there was a diversity in the length of putative uORF genes (Fig 5B and Supplementary Table S11). Interestingly, most of the predicted amino acid sequences showed no homology to each other, while those with detectable similarity could be classified into four distinct groups, hereafter referred to as uORF types A–D (Fig 5C and Supplementary Table S12). The putative uORF of the AUS camel picornavirus 1 virus belongs to uORF type A.

**Fig 5.**
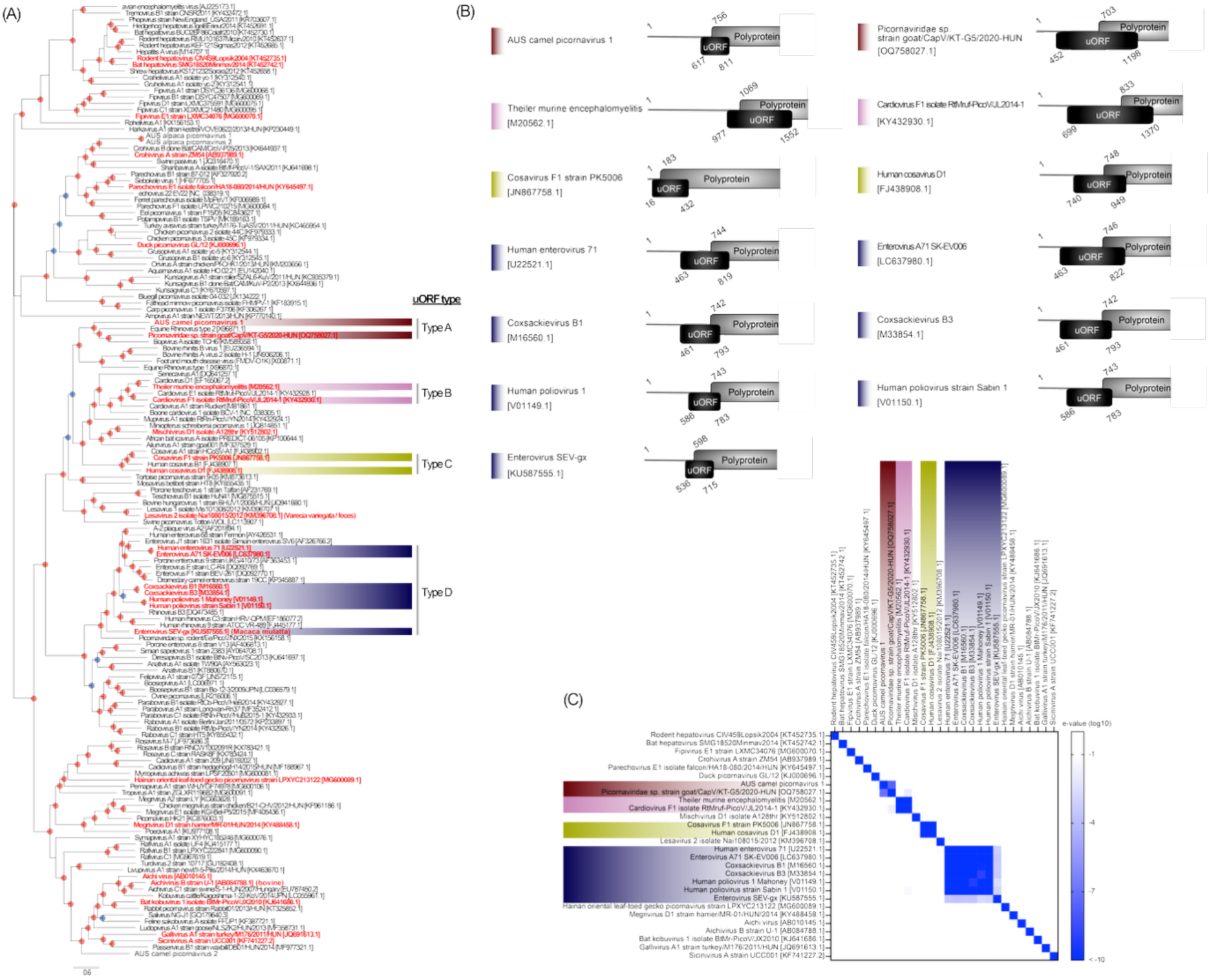
Phylogenetic relationships and homology of putative uORFs in AUS camel picornavirus 1 and other members of the *Picornaviridae*. (A) Maximum likelihood phylogenetic tree inferred from amino acid sequences of the picornavirus polyprotein, including viruses identified in this study and representative reference viruses. Branch lengths represent the number of substitutions per site. Red and blue circles at internal nodes indicate bootstrap values ≥ 90% and ≥ 80%, respectively. The tree was midpoint rooted for clarity only. Viruses with (red) or without (black) putative uORFs are indicated. Colours denote uORF types: dark red represents type A, pink represents type B, yellow represents type C, and blue represents type D. (B) Schematic overview of the putative uORF in AUS camel picornavirus 1 and representative viruses. (C) Homology of each putative uORF-encoded protein. Pairwise E-values of putative uORF-encoded proteins were calculated using BLASTP against a custom database comprising the same set of uORF- encoded protein sequences used as queries, thereby enabling direct pairwise comparisons within this sequence group. For cases with multiple hits on the same protein, the hit with the longest alignment length was used. Hits with log10(E-value) ≥ –1 were considered as having no homology. The full matrix of pairwise E-values is provided in Supplementary Table S12.

### Structural and functional similarity in uORF-encoded proteins across multiple lineages

To assess structural similarity among uORF-encoded proteins, we focused on secondary structure and intrinsic disorder profiles rather than three-dimensional fold homology as short and rapidly evolving proteins frequently lack stable tertiary structures and hence are often better described in terms of their secondary structure tendencies and intrinsic disorder rather than well-defined folds [18, 19]. Using PSIPRED [20, 21] and AIUPred predictions [22], uORF-encoded proteins were classified as ordered-helix-rich when the fraction of residues predicted to form ordered α-helices was ≥ 0.3, indicating a substantial proportion of residues predicted to form ordered α-helical structures. Despite limited amino acid sequence similarity among uORF-encoded proteins identified in phylogenetically independent viral lineages, a large subset of uORF-encoded proteins met this criterion (n = 20/28, 71.4%), revealing a recurrent structural propensity (Fig 6A and Supplementary Table S13). Mapping this structural classification onto the viral phylogeny showed that ordered-helix-rich uORF-encoded proteins were present in multiple independent lineages (Fig 6B), suggesting the repeated emergence of broadly comparable secondary structure composition and intrinsic disorder patterns. These findings indicate that although uORF-encoded proteins lack strong primary sequence conservation, they repeatedly evolve comparable structural tendencies across divergent viral lineages.

**Fig 6.**
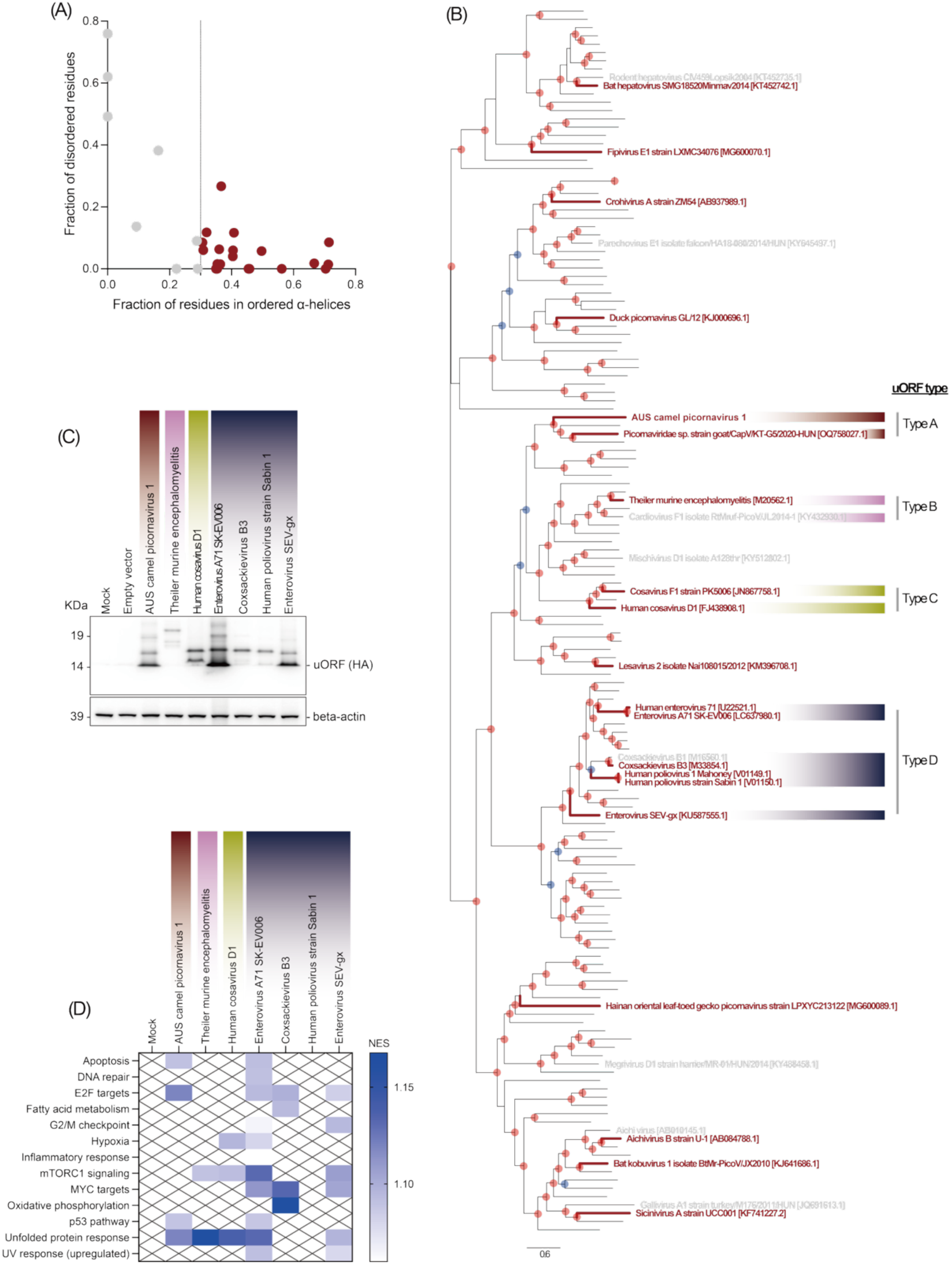
Structural comparison and functional assessment of uORFs across diverse picornaviruses. (A) Scatter plot showing the relationship between ordered helix fraction and disorder fraction for uORF-encoded proteins. Each point represents a single uORF-encoded protein; burgundy points indicate ordered-helix-rich uORF-encoded proteins (fraction of residues in ordered α-helices ≥ 0.3), grey points indicate all other uORF-encoded proteins, and the vertical dashed line denotes the threshold used to define ordered-helix-rich uORF-encoded proteins. (B) Mapping of ordered-helix-rich uORF-encoded proteins onto the viral phylogeny. Tip labels are coloured according to structural classification, with ordered-helix-rich uORF-encoded proteins (fraction of residues in ordered α-helices ≥ 0.3) shown in burgundy and all other uORF-encoded proteins shown in grey. The phylogeny is identical to that shown in Fig 5A. Tip labels for viruses that do not contain an uORF have been removed. Colours denote uORF types: dark red represents type A, pink represents type B, yellow represents type C, and blue represents type D. (C) Western blot analysis of 3×HA-tagged uORF-encoded proteins expressed in HEK293T cells. Cell lysates were probed with an anti-HA antibody to detect uORF-derived proteins, and β-actin was detected as a loading control. (D) Pathway-level transcriptional responses induced by distinct uORFs. Heatmap showing normalized enrichment scores (NES) for selected Hallmark gene sets derived from Gene set enrichment analysis (GSEA) of RNA-seq data. For each uORF, gene expression profiles were compared with the 3×HA-tagged empty-vector control, and non-significant pathways (adjusted *p*-value ≥ 0.1) were masked. Differential expression analysis was performed using three independent biological replicates per condition (n = 3). Gene-level fold changes and *p*-values from this analysis are provided in Supplementary Table S14 and were used to generate ranked gene lists for fast gene set enrichment analysis (fgsea). The complete fgsea results for all Hallmark gene sets across all comparisons are provided in Supplementary Table S15.

To investigate whether the putative uORF of the diverse viruses in the *Picornaviridae,* including AUS camel picornavirus 1, encodes a functional protein and shares functional similarities, we cloned each uORF into an expression vector with a C-terminal 3×HA tag and performed gene expression analysis of cells expressing these constructs. First, we confirmed stable expression of the uORF-derived proteins by western blotting (Fig 6C). For all uORFs, bands of the sizes expected based on their predicted lengths were consistently detected, indicating successful expression of the encoded protein products. In addition, higher-molecular-weight species were observed for each uORF as a recurring feature. Subsequently, we performed transcriptomic analysis by transfecting HEK293T cells with each uORF construct and comparing gene expression profiles to empty-vector controls. A non-transfected mock condition was included to account for transfection-independent effects. Although individual uORFs induced largely distinct gene-level transcriptional changes (Supplementary Fig S4 and Supplementary Table S14), pathway-level enrichment analysis using GSEA revealed partially overlapping patterns of pathway activation across multiple uORFs. In particular, subsets of uORFs reproducibly enriched related host pathways, including the unfolded protein response, mTORC1 signaling, and cell cycle-associated gene sets (e.g., E2F and MYC targets) (Fig 6D), although no single pathway was uniformly affected across all uORFs. In contrast, these pathways were not significantly enriched in mock-transfected controls (Fig 6D). These results suggest that despite minimal sequence similarity, distinct uORFs can reproducibly modulate overlapping host pathways.

## Discussion

The introduction of non-native animal species into the geographically isolated Australian continent had major ecological consequences for the resident flora and fauna and may also have impacted RNA virus evolution. In particular, invasive animals (e.g., camels in Australia) may introduce new RNA viruses or different genotypes of known RNA viruses into the Australian ecosystem, from where they may jump species boundaries to emerge in new hosts. Previous work on Australian camels has been limited to serological surveillance for Middle East respiratory syndrome coronavirus (MERS-CoV) [23], for which no evidence has been found. To explore viral diversity more broadly, we conducted virome analyses of camelid populations from different regions of Australia. As in earlier studies, we found no evidence of MERS-CoV. Notably, however, we did identify distinct picornaviruses in Australian camelids that are related to viruses previously detected in camels or other artiodactyls from geographic locations distant from Australia. This phylogenetic pattern raises the possibility that these viruses were introduced with the founding camel populations and subsequently persisted and diversified in Australia.

One of the most important questions in the biology of invasive species is whether they transmit their viruses to the indigenous native species, or if they primarily acquire viruses from the resident host taxa. With respect to the camelids studied here, neither of these simple scenarios adequately explains the observed patterns. Notably, we did not detect viruses in these camelids that were closely related to those previously described in Australian native or domestic animals, nor did we observe the phylogenetic intermixing among hosts that is indicative of recent cross-species transmission within Australia. However, this result should be interpreted with caution because of the limited virome data currently available from native species that share ecological niches with camelids in Australia. Notably, the astrovirus and picornaviruses described here were detected in camelid populations from Western Australia and New South Wales, regions where ecological contact with domestic livestock and native wildlife is plausible, rather than in geographically remote populations in central Australia. Despite these potential transmission opportunities, we found no evidence of recent phylogenetic intermixing with viruses from Australian domestic or native hosts. More comprehensive sampling of native host viromes is required to test this hypothesis. Rather than evidence for ongoing viral exchange between invasive and native hosts in Australia, the viromes we describe are more consistent with founder effects at the time of introduction, followed by diversification within the invasive host populations themselves, leading to the evolution of unique viral lineages. Hence, even when invasive species have experienced major population expansions that might be expected to promote frequent cross-species transmission, the dominant signal in viral evolutionary history reflects a deeper evolutionary history and population divergence.

Of note, we detected a virus closely related to avian astroviruses in Australian camels. This phylogenetic pattern – of a mammalian virus clustering with viruses previously found in birds – strongly suggests that an avian-derived astrovirus was transmitted across vertebrate classes to camels. Astroviruses infect a wide range of vertebrates, and are broadly classified into “mamastroviruses” that infect mammals and “avastroviruses” that infect birds, and these two group fall as separate lineages in phylogenetic analyses [24, 25]. Such a phylogenetic pattern indicates that astrovirus evolution has proceeded based on relatively strong host specificity. Despite this, viruses related to avian astroviruses have been found in rodents, and astroviruses related to rodents have been found in birds [26]. These findings suggest that the evolutionary divergence between avian and mammalian astroviruses is less strict than previously thought. In addition, that the avastroviruses fall within the diversity of the mamastroviruses (unless phylogenies are explicitly rooted on the avastrovirus-mamastrovirus split), indicates a longer evolutionary history of cross-species transmission events. However, the limited sampling of camelids currently precludes inference of whether this host-jump occurred in Australia or elsewhere, highlighting the need for broader global sampling of camels.

The discovery of a putative uORF within a highly divergent picornavirus lineage, together with comparable elements identified across other taxa in the family, suggests that the acquisition of novel coding sequences upstream of the polyprotein ORF may be a recurrent feature of picornavirus evolution. Although most of these uORF-encoded proteins exhibit no amino acid sequence homology, overlapping secondary structure composition and shared functional effects indicate that similar biological outcomes could sometimes be achieved through distinct sequence solutions. This pattern of no sequence homology coupled with analogous functional outputs is consistent with an evolutionary process in which uORFs are repeatedly gained or lost depending on selective pressures acting at the host–virus interface. The multiple independent events of uORF gain and loss across the phylogeny further suggest that these elements are unlikely to derive from a single ancestral innovation, but instead may arise through recurrent cycles of origination and loss of small accessory proteins. Consistent with this idea, recent studies across multiple RNA virus families, including the *Coronaviridae*, *Filoviridae*, *Flaviviridae*, *Retroviridae*, and *Picornaviridae*, have demonstrated that uORFs can modulate viral replication, tissue tropism, and interactions with the host immune system [11–16]. Importantly, the presence of uORFs in distinct viral lineages implies that their contribution to viral fitness is context-dependent rather than universally required. In addition, a previous study of human enteroviruses illustrated that recombination frequently occurs at the 5′UTR–capsid boundary, a region regarded as a major recombination “hotspot” within the genus [27]. These findings reinforce the view that upstream genomic regions act as evolutionary “hotspots” in RNA virus genomes, providing a permissive landscape for the repeated emergence of small accessory ORFs. Notably, within the *Picornaviridae*, we found evidence that independently acquired uORFs can converge on similar functional behaviours when occupying equivalent genomic locations, suggesting that lineage-specific constraints and recurrent selective pressures favour analogous functional solutions.

The presence or absence of uORFs in picornaviruses varies among lineages that occupy distinct host environments. For example, several respiratory-tropic picornaviruses, including enterovirus D68 and rhinoviruses [13], lack identifiable uORFs, and the equine rhinitis B virus—closely related to the lineage reported here—similarly does not encode this feature. In contrast, uORFs are found in multiple viral lineages that infect the gastrointestinal tract, and the representative uORF-encoded proteins examined here were associated with partially overlapping pathway-level transcriptional responses when expressed in host cells. Most notably, uORFs from several distinct lineages, including AUS camel picornavirus 1, Theiler murine encephalomyelitis, Human cosavirus D1, Enterovirus A71 SK-EV006, and Enterovirus SEV-gx viruses, consistently showed positive enrichment of the unfolded protein response (UPR) pathway. In addition, enrichment of mTORC1 signalling was observed across multiple uORFs, including those from Theiler murine encephalomyelitis, Human cosavirus D1, Enterovirus A71 SK-EV006, and Enterovirus SEV-gx viruses, suggesting recurrent modulation of host translational and metabolic pathways. A third common feature was the enrichment of cell cycle–associated gene sets, such as E2F and MYC targets, which was observed for uORFs from AUS camel picornavirus 1, Enterovirus A71 SK-EV006, Coxsackievirus B3, and Enterovirus SEV-gx viruses. Together, these results indicate that distinct uORFs repeatedly target a limited set of host pathways related to protein homeostasis, translation, and cell cycle regulation.

These observations raise the possibility that uORFs function as accessory modules that depend on tissue-specific physiological conditions. The intestinal environment is characterized by high protein synthesis demands, rapid epithelial turnover, and tightly regulated proliferative programs, which may favour viral strategies that modulate UPR, mTORC1 signalling, and cell cycle–associated pathways [28–31]. Notably, the uORF-encoded proteins analysed here share little or no detectable amino acid sequence similarity and do not induce identical transcriptional programs, and the observed partial and pathway-restricted overlap is consistent with multiple independent acquisitions shaped by convergent selective pressures acting on a shared set of host regulatory pathways. In enteroviruses, uORFs have been experimentally shown to influence viral replication efficiency in differentiated human intestinal organoids [13], highlighting their potential functional relevance in specific cellular contexts. However, because infectious AUS picornavirus 1 particles were not isolated in this study, the effects of the identified uORFs on viral replication efficiency, as well as their expression levels in infected cells, remain unknown. Accordingly, additional experimental studies are needed to directly evaluate the functional roles of these uORFs in the context of viral infection.

Collectively, our study reveals that invasive species can harbour genetically divergent viruses that are absent from native species, including highly divergent picornavirus lineages. We further suggest that uORFs represent a dynamic genomic feature subject to recurrent origination and loss, and are potentially maintained in specific ecological or cellular contexts. Experimental analyses revealed that uORFs from divergent lineages can elicit similar transcriptional responses in host cells, pointing to context-dependent convergence in functional outputs.

## Materials and Methods

### Ethical approval and scientific licenses

The collection of samples from alpacas, camels and llamas was approved by Taronga Conservation Society Australia’s Animal Ethics Committee (approval number 4c/10/21) and the University of Sydney Animal Ethics Committee (approval number 2022/2096).

### Sample collection and processing

Faeces, rectal and nasal swabs, saliva, eye swabs, and milk were collected between January 2022 and May 2023 from alpacas (*Vicugna pacos*), dromedary camels (*Camelus dromedarius*), and llamas (*Lama glama*) across Australia. The majority (75%) were faecal samples obtained from multiple locations across three Australian states: Sutton, Muswellbrook, Anna Bay, Dubbo, and Mosman (New South Wales, NSW); Yulara (Northern Territory, NT); and Hopeland, Perth, Kordabup, Morangup, Esperance, Youngs Siding, Myrup, and Rosa Glen (Western Australia, WA). Nasal swabs and saliva samples were collected from selected locations in two states: Hopeland and Youngs Siding in WA, and Muswellbrook and Anna Bay in NSW. All samples were preserved in RNAlater and stored at –80°C, except for milk samples.

### Sample processing and sequencing

Individual aliquots of each sample were mixed with 600 μl of lysis buffer containing 1% β-mercaptoethanol (Sigma-Aldrich). Faecal material and swab samples were homogenized using QIAshredder columns (Qiagen), and total RNA was extracted from the resulting supernatants using the RNeasy Plus Mini Kit (Qiagen), according to the manufacturer’s instructions. Extracted RNA was then pooled in approximately equimolar amounts, and grouped by location, sample type, and animal species. This yielded 56 RNA pools, each comprising a median of six samples (range: 1–9) (Supplementary Tables S1). Sequencing libraries were prepared using the TruSeq Total RNA Library Preparation Protocol (Illumina), with host ribosomal RNA removed via the Ribo-Zero Plus Kit (Illumina). Paired-end sequencing (150 bp) was conducted on the NovaSeq 6000 platform (Illumina) at the Australian Genome Research Facility (AGRF) in Melbourne, Australia. Libraries generated from each lane were processed and analysed independently. For genome assembly and abundance estimation, read data from both lanes were combined and treated as a single data set.

### Identification of novel virus sequences

Sequencing libraries were processed using the BatchArtemisSRAMiner pipeline (v1.0.4) [32]. Briefly, raw reads were quality-trimmed and adapter sequences removed with Trimmomatic (v0.38) [33] using the parameters SLIDINGWINDOW:4:5, LEADING:5, TRAILING:5, and MINLEN:25, prior to assembly. *De novo* assembly was performed with MEGAHIT (v1.2.9) [34]. The resulting contigs identified were then screened against the RdRp-scan core RNA-dependent RNA polymerase (RdRp) protein database (v0.90) [35] and the protein version of the Reference Viral Databases (RVDB v26.0) [36, 37] using DIAMOND BLASTx (v2.1.6) [38]. An e-value cut-off of 1 × 10⁻⁴ was applied for RdRp-scan and 1 × 10⁻¹⁰ for RVDB. To reduce false positives, contigs with viral hits were further queried against the NCBI nucleotide database (as of July 2024) using BLASTN [39], and against the NCBI non-redundant protein (nr) database using DIAMOND BLASTx. Viral candidates were evaluated based on sequence similarity to known vertebrate viruses (e.g., percentage amino acid identity) and, in some cases, phylogenetic placement. Sequences likely originating from non-vertebrate viruses (such that they were likely of dietary origin) were excluded from downstream analyses.

To determine whether a sequence represented a novel virus, we followed the species demarcation criteria established by the International Committee on Taxonomy of Viruses (ICTV) for each virus family. In the absence of formal criteria, sequences were considered novel if they exhibited less than 95% amino acid identity to known viruses across the RdRp region and formed a distinct phylogenetic group.

Polyprotein annotation for viruses in the *Picornaviridae* was performed using BLASTp (v2.15.0), and the genomic positions of BLAST hits were used to infer the approximate boundaries of structural (VP0–VP1) and non-structural (2A–3D) proteins. Crohivirus A (NC_025474.1), Equine rhinitis B virus 1 (NC_003983.1), and Caprine kobuvirus sequence 12Q108 (NC_023422.1) were used as reference genomes for AUS alpaca picornavirus 1/2, AUS camel picornavirus 1, and AUS camel picornavirus 2, respectively. Conserved motifs were inspected to refine protein boundaries, and candidate 3C cysteine protease cleavage sites were identified by scanning for common picornaviral motifs (i.e., QG, QS, QA, QV, and EG) [40, 41] (Supplementary Tables S7-S9). uORFs were predicted in Geneious Prime (v 2025.1.3) using the built-in ORF finder, and were defined as upstream open reading frames longer than 150 nucleotides located in the 5′ region preceding the polyprotein ORF, initiated by an AUG start codon, including those overlapping the polyprotein initiation codon.

### Genome abundance

The relative abundance of each viral contig was calculated using the expected count values generated by RNA-Seq by the Expectation Maximization (RSEM) software (v1.3.0)[42]. Abundance values were expressed as the proportion of reads mapped to each viral contig relative to the total number of trimmed reads in the corresponding library.

### Phylogenetic analysis

Amino acid sequences of each relevant viral group were aligned using the L-INS-i protocol in MAFFT version 7.453 [43]. Ambiguously aligned regions were trimmed using trimAl v1.5 [44], with a gap threshold of 0.9 and a minimum conservation threshold of 60%. Maximum likelihood phylogenetic trees were then inferred using IQ-TREE 2 v2.3.6 [45], with the optimal substitution model selected by ModelFinder [46]. Branch support was estimated using 1,000 bootstrap replicates with the UFBoot2 algorithm and an implementation of the SH-like approximate likelihood ratio test [47] available within IQ-TREE 2.

### Assessment of sequencing library composition

To verify the composition of potential host species present in libraries where camel astrovirus was detected (see Results), taxonomic composition was assessed using Kraken2 (v2.1.2) [48, 49]. Accordingly, Kraken2 was run on trimmed reads with the non-default options --minimum-hit-groups 3 and --report-minimizer-data, using the nt database from May 30, 2024 (available at https://benlangmead.github.io/aws-indexes/k2). To estimate species-level abundances, Bracken (v2.9) [50] was applied to the Kraken2 output, and the results were processed and visualized using KrakenTools (v1.2) [51] and Krona (v2.8.1) [52].

### Secondary structure and intrinsic disorder analysis of uORF-encoded proteins

Amino acid sequences of predicted uORF-encoded proteins were analysed for secondary structure and intrinsic disorder. Secondary structure was predicted using the PSIPRED web server (default parameters) [20, 21], which assigns each residue to α-helix, β-strand, or coil. Intrinsic disorder was predicted using the AIUPred web server (default parameters) [22], which provides per-residue disorder scores ranging from 0 to 1. Residues with disorder scores ≥ 0.5 were classified as disordered. For each uORF-encoded protein, residue-level predictions were summarized into uORF-level metrics. The fraction of residues predicted as α-helix (helix fraction), β-strand, and coil was calculated relative to uORF-encoded protein length. The fraction of disordered residues (disorder fraction) was calculated as the proportion of residues with AIUPred scores ≥ 0.5. To integrate secondary structure and disorder predictions, α-helical residues were further subdivided into ordered and disordered helices: the ordered helix fraction was defined as the proportion of residues predicted as helix with disorder scores < 0.5, whereas the disordered helix fraction was defined as the proportion of residues predicted as helix with disorder scores ≥ 0.5. α-helical content was used as a descriptive metric of structural composition and as a pragmatic proxy for potential interaction interfaces, given that short α-helical elements can contribute to molecular recognition in small proteins and peptides [19]. uORF-encoded proteins were classified as ordered-helix-rich when ≥30% of residues were predicted to form ordered α-helices. This threshold was defined empirically based on the distribution of ordered helix fractions across all uORF-encoded proteins (Fig. 6A), which revealed a subset of proteins with relatively elevated ordered helical content. All uORF-level metrics were compiled into a single table for downstream comparative analyses. Internal consistency was verified by confirming that the helix fraction equalled the sum of ordered and disordered helix fractions, with only negligible numerical deviations attributable to floating-point rounding.

### Cells

HEK293T cells (obtained from the National Institute of Infectious Disease, Tokyo, Japan) were maintained in Dulbecco’s Modified Eagle Medium (DMEM; High Glucose) with L-Glutamine and Phenol Red (Wako, 044–29765) supplemented with 10% FBS and 1% Penicillin-Streptomycin Solution. The cells were maintained at 37 °C with 5% CO2.

### Western blotting and transcriptomic analysis

The upstream open reading frame (uORF) gene of the camel erbovirus-like virus, Theiler murine encephalomyelitis (GenBank accession M20562.1), Human cosavirus D1(GenBank accession FJ438908.1), Enterovirus A71 SK-EV006 (GenBank accession LC637980.1), Coxsackievirus B3 (GenBank accession M33854.1), Human poliovirus Sabin 1(GenBank accession V01150.1) and Enterovirus SEV-gx (GenBank accession KU587555.1) were cloned into pCAGGS vector with a C-terminal 3×HA tag. HEK293T cells were transfected with the uORF-encoded protein expression plasmids using TransIT-LT1 (Takara, Cat# MIR2305). At 24 hours post-transfection (hpt), cells were lysed in buffer containing 20 mM Tris-HCl (pH 7.4), 135 mM NaCl, 1% Triton X-100, 1% glycerol, and protease inhibitor tablets (Roche). Lysates were incubated on ice for 20 min and clarified by centrifugation at 15,000 rpm for 5 min at 4°C. Supernatants were mixed with an equal volume of 2× SDS sample buffer (50 mM Tris-HCl, pH 6.8, 4% SDS, 0.2% bromophenol blue, 10% glycerol, 200 mM β-mercaptoethanol) and heated at 95°C for 5 min. Proteins were separated by SDS-PAGE (NuPAGE, Life Technologies) and transferred onto nitrocellulose membranes (iBlot2, Life Technologies). Membranes were blocked in PBS containing 0.05% Tween-20 (PBS-T) and 5% skim milk for 1 hour (h) at room temperature, and then incubated at 4°C for 24 h with primary antibodies: anti-HA tag (mouse IgG, Cat# M180-3; MBL Life science) or anti-β-actin (rabbit IgG, Cat# A2228; Merck). After three washes with PBS-T, membranes were incubated for 1 h at room temperature with HRP-conjugated goat anti-mouse or goat anti-rabbit IgG secondary antibodies, followed by three additional washes with PBS-T. Antibody dilutions were performed according to the manufacturer’s instructions. Signals were detected using Amersham ECL Western Blotting Detection Reagents (GE Healthcare), and imaged with the Amersham Imager (GE Healthcare).

### Transcriptome data processing and differential gene expression analysis

Total RNA was extracted from HEK293T cells from three independent biological replicates per condition at 24 hpt using the ISOGEN II (Nippon Gene) according to the manufacturer’s protocol. Sequencing libraries were prepared from the RNAs using a TruSeq stranded mRNA sample prep kit (Illumina, San Diego, CA) according to the manufacturer’s protocol. The library was sequenced using an Illumina NovaSeq 6000 platform in a 100 bp single-end mode. Raw sequencing reads were subjected to quality trimming using Trimmomatic [53] v0.39 to remove adaptor sequences and low-quality bases. The trimmed reads were then aligned to the human reference genome (GRCh38) using HISAT2 [54] v2.1.0 with default parameters. Gene-level read counts were generated using featureCounts [55] v2.0.6 based on annotated gene models. Variance-stabilized gene expression values were used for exploratory analyses, including principal component analysis (Supplementary Fig S4). Differential gene expression analysis was performed using Cuffdiff [56] v2.2.1 by comparing each uORF-expressing condition to the empty-vector control. For each comparison, fold changes and associated *p*-values were obtained for all detected genes (Supplementary Table S14). No minimum fold-change or significance threshold was applied prior to downstream pathway analysis. Summary statistics of sequencing depth and mapping efficiency for all RNA-seq samples, including total read counts and mapping rates, are provided in Supplementary Table S16.

### Pathway enrichment analysis

For pathway-level analysis, Gene Set Enrichment Analysis (GSEA) [57] was conducted using ranked gene lists derived from differential expression results. Genes were ranked according to a signed significance metric calculated as the sign of the log_2_ fold change multiplied by the negative log_10_-transformed *p*-value. Enrichment analysis was performed using the fgsea package with the Hallmark gene set collection from MSigDB [58, 59]. Gene sets containing fewer than 15 or more than 500 genes were excluded. Statistical significance was assessed using permutation testing, and multiple testing correction was applied using the Benjamini–Hochberg method [60]. Pathways with an adjusted *p*-value < 0.1 were considered significant.

## Acknowledgments

This work was funded by a National Health and Medical Research Council (NHMRC) Investigator award (GNT201719) to E.C.H, and by a KAKENHI Grant-in-Aid for Early-Career Scientists (25K18814) to K.T and RIKAKEN HOLDINGS CO. Young Researcher Support Grant-in-aid to K.T.

We thank Dr. Karrie Rose of the Australian Registry of Wildlife Health, Taronga Conservation Society Australia for generously providing samples and for valuable discussions. Taronga Conservation Society Australia is also acknowledged for their commitment to wildlife health investigation through long-term support of the Australian Registry of Wildlife Health. We are grateful to the farm owners and staff for their assistance with sample collection.Data analysis was conducted using the software available on the University of Sydney HPC Service (Artemis) and the Setonix Supercomputer at the Pawsey Supercomputing Research Centre, Australia. The transcriptomic profiling of cultured cells was performed with sequencing support from the NGS Core Laboratory, Genome Information Research Center, Institute for Microbial Diseases, the University of Osaka.

## Author contributions

K.T., J.C.O.M., and E.C.H. conceptualized this study. J.C.O.M., E.H. and S.S. performed the animal sampling. B.J.L. processed the samples. K.T. and J.C.O.M. analysed data. K.T., J.H., and Y.M. performed the protein experiments and/or interpreted the results. K.T. wrote the original draft. J.C.O.M., J.H and E.C.H contributed to manuscript revisions and additions. All authors reviewed and approved the final manuscript.

## Conflict of interest

The authors declare no conflicts of interest.

## Data availability

Raw read sequences generated for this project have been deposited at the Sequence Read Archive (SRA) database under Bioproject: PRJNA1423894 (BioSample accessions SAMN55370158 – SAMN55370325) and PRJNA1419329 (BioSample accessions SAMN55129586 – SAMN55129612), and GenBank database (accession numbers: PX987501– PX987505).

## Notes

### Competing Interest Statement

The authors have declared no competing interest.

