## Supplementary Information for "RNA virus discovery in Australian camelids reveals divergent picornaviruses and the convergent evolution of upstream ORFs"

- Supplementary results
- Supplementary Figures S1–S4
- Supplementary Tables S1–S16 (separate Excel files)

### Supplementary results

#### Individual gene phylogenies of novel viruses.

To more precisely document the evolutionary position of AUS alpaca picornavirus 1 and alpaca picornavirus 2, we performed phylogenetic analyses on individual amino acid alignments of the 3CD protein (i.e., the precursor comprising 3C cysteine protease and 3D RdRp), 3D protein (i.e., RdRp) and P1 region (encoding VP0, VP3 and VP1), together with representative related viruses (i.e., genus *Crohivirus*). To achieve this, we first identified the genomic positions of structural (i.e., P1 region) and non-structural (3CD and 3D) proteins (Supplementary Fig S2A and Supplementary Table S7). Phylogenetic analyses yielded consistent but lineage-specific patterns (Supplementary Fig S2B–S2D). The 3D protein of AUS alpaca picornavirus 1 and alpaca picornavirus 2 showed the highest amino acid sequence identity (52.93%) to Crohivirus A [NC_025474.1]. Taken together, these data confirmed that AUS alpaca picornavirus 1 and AUS alpaca picornavirus 2 belong to the same species and form a genetically distinct lineage that is sufficiently genetically divergent to justify classification as a novel species within the genus *Crohivirus* (family *Picornaviridae*).

Using the same analytical framework, we examined the evolutionary position of the AUS camel picornavirus 2 sequence. Individual amino acid alignments of the 3CD, 3D, and structural proteins (i.e., P1 region) were constructed with representative viruses from the genera *Kobuvirus*, *Salivirus*, *Sakobuvirus*, *Ludopivirus*, *Gallivirus*, *Passerivirus*, and *Sicinivirus*. Protein boundaries were inferred as described above (Supplementary Fig S3A and Supplementary Table S8). Phylogenetic analyses of these sequence alignments revealed that each AUS camel picornavirus 2 protein formed a well-supported cluster that included closely related sequences previously detected from Bactrian camels in China (Supplementary Fig S3B–S3D). In the 3D phylogeny, the AUS camel picornavirus 2 sequence grouped with *Kobuvirus* and *Salivirus* (Supplementary Fig 3C). In contrast, in the P1 and 3CD phylogenies it was positioned outside the established genera *Kobuvirus*, *Salivirus*, *Sakobuvirus*, *Ludopivirus*, *Gallivirus*, *Passerivirus*, and *Sicinivirus*, forming a distinct lineage within a broader clade (Supplementary Fig S3B and S3D). The 3D sequence of AUS camel picornavirus 2 showed the highest amino acid sequence identity (94.08%) to *Picornaviridae* sp. QXDL/86006 (PX113272.1) detected in bactrian camels in China. Taken together, these data suggest that AUS camel picornavirus 2 likely represents a camel-specific lineage that is sufficiently genetically distinct to warrant classification as novel genus within the *Picornaviridae*.

### Supplementary Figures

**Supplementary Figure S1. Krona chart of eukaryotic reads detected in camelid samples.** Reads were first classified into Bacteria, Archaea, Eukaryota, and Viruses and unclassified sequences, and their relative proportions are shown in the first pie chart. The second chart shows the composition of Eukaryota reads, including Mammalia, Aves, and other eukaryotic groups (such as Mastigamoebida, Insecta, and Litostomatea), with Mammalia further divided into Artiodactyla and non-Artiodactyla. Percentages in parentheses indicate the proportion of each category relative to the total reads.

**Supplementary Figure S2. Individual gene phylogenies of AUS alpaca picornavirus 1 and 2.**

(A) Schematic overview of structural (P1 region) and non-structural (2A–3D) genes in AUS alpaca picornavirus 1 and 2. (B-D) Maximum likelihood phylogenetic trees based on amino acid sequences of the (B) 3CD and (C) 3D (RdRp) proteins, and (D) P1 region, including the AUS alpaca picornavirus 1 and 2 as well as representative close-related sequences. Branch lengths represent the number of substitutions per site. Red and blue circles at internal nodes indicate bootstrap values ≥ 90% and ≥ 80%, respectively. Based on their phylogenetic positions inferred from the polyprotein tree (Fig 3), Parechovirus sp. PPYCC16/10 (OR867093.1) and Chalinolobus tuberculatus picornavirus B10 (OR248863.1) were used as outgroups for the corresponding 3CD, 3D, and P1 phylogenetic analyses, respectively.

**Supplementary Figure S3. Individual gene phylogenies of AUS camel picornavirus 2.**

(A) Schematic overview of structural (P1 region) and non-structural (2A–3D) genes in AUS camel picornavirus 2. (B-D) Maximum likelihood phylogenetic trees based on amino acid sequences of the (B) 3CD and (C) 3D (RdRp) proteins, and (D) P1 region, including the AUS camel picornavirus 2 and representative closely related sequences. Sequences derived from camels outside Australia are highlighted in brown. Branch lengths represent the number of substitutions per site. Red and blue circles at internal nodes indicate bootstrap values ≥ 90% and ≥ 80%, respectively. Based on their phylogenetic positions inferred from the polyprotein tree (Fig 3), Turdivirus 2 10717 (GU182408.1) and Livupivirus A1 newt/II-5-Pilis/2014/HUN (KX463670.1) were used as outgroups for the corresponding 3CD, 3D, and P1 phylogenetic analyses, respectively.

**Supplementary Figure S4. Principal component analysis (PCA) of global gene expression profiles in HEK293T cells transfected with each 3×HA-tagged uORF expression vector, 3×HA-tagged empty vector, or mock-treated controls.**
PCA of variance-stabilized gene expression values from HEK293T cells transfected with 3×HA-tagged uORF expression vectors, 3×HA-tagged empty vector, or mock-treated controls was performed as an exploratory analysis. PC1–PC2 and PC1–PC3 projections are shown, with each point representing an independent biological replicate.

### Supplementary Tables S1-S16

**Supplementary Table S1.** Samples used for RNA sequencing.

**Supplementary Table S2.** Predicted protein domains identified in AUS camel astrovirus genomes using BLASTX.

**Supplementary Table S3.** Predicted protein domains identified in AUS alpaca picornavirus 1 genomes using BLASTX.

**Supplementary Table S4.** Predicted protein domains identified in AUS alpaca picornavirus 2 genomes using BLASTX.

**Supplementary Table S5.** Predicted protein domains identified in AUS camel picornavirus 1 genomes using BLASTX.

**Supplementary Table S6.** Predicted protein domains identified in AUS camel picornavirus 2 genomes using BLASTX.

**Supplementary Table S7.** Predicted non-structural and structural proteins encoded by the polyprotein in AUS alpaca picornavirus 1 and 2.

**Supplementary Table S8.** Predicted non-structural and structural proteins encoded by the polyprotein in AUS camel picornavirus 1.

**Supplementary Table S9.** Predicted non-structural and structural proteins encoded by the polyprotein in AUS camel picornavirus 2

**Supplementary Table S10.** Summary of complete non-redundant picornavirus genomes containing putative upstream ORFs (uORFs), including *Erbovirus*, *Bopivirus*, *Aphthovirus*, and related unclassified lineages closely related to the AUS camel picornavirus 1.

**Supplementary Table S11.** Summary of complete non-redundant picornavirus genomes containing putative upstream ORFs (uORFs) identified from NCBI Virus.

**Supplementary Table S12.** Pairwise BLASTP E-values among putative uORF-encoded proteins.

**Supplementary Table S13.** Secondary structure and intrinsic disorder metrics of uORF-encoded proteins.

**Supplementary Table S14.** Gene-level differential expression statistics used for fgsea in HEK293T cells transfected with each 3×HA-tagged uORF expression vector or mock-treated controls, compared with 3×HA-tagged empty vector controls.

**Supplementary Table S15.** Complete fgsea results for all Hallmark gene sets across all comparisons in HEK293T cells transfected with each 3×HA-tagged uORF expression vector or mock-treated controls, compared with 3×HA-tagged empty vector controls.

**Supplementary Table S16.** RNA-seq read counts and mapping statistics for all HEK293T cell samples transfected with each 3×HA-tagged uORF expression vector, a 3×HA-tagged empty vector, or mock controls.
