## Supplementary figures and images for "RNA virus discovery in Australian camelids reveals divergent picornaviruses and the convergent evolution of upstream ORFs"

### Supplementary Figure S1

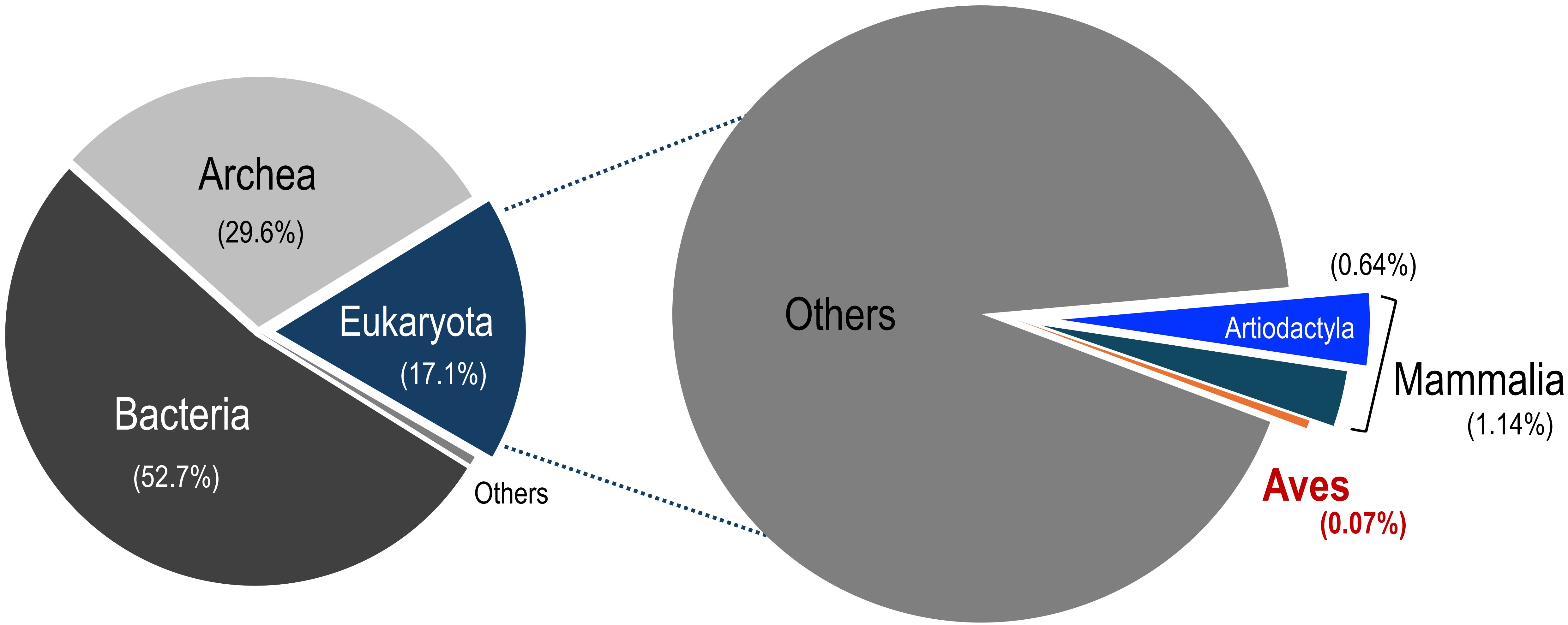

### Supplementary Figure S2

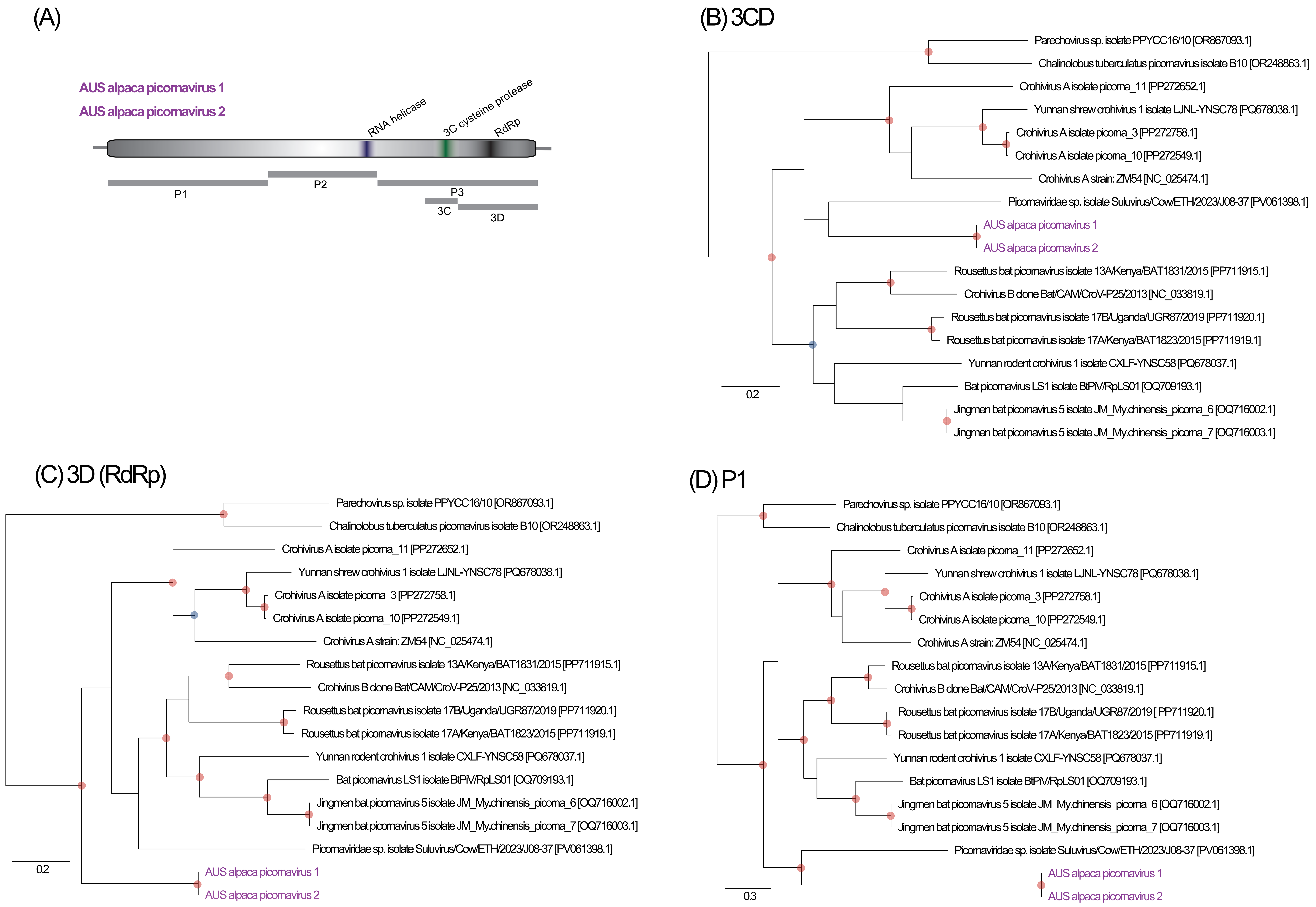

### Supplementary Figure S3

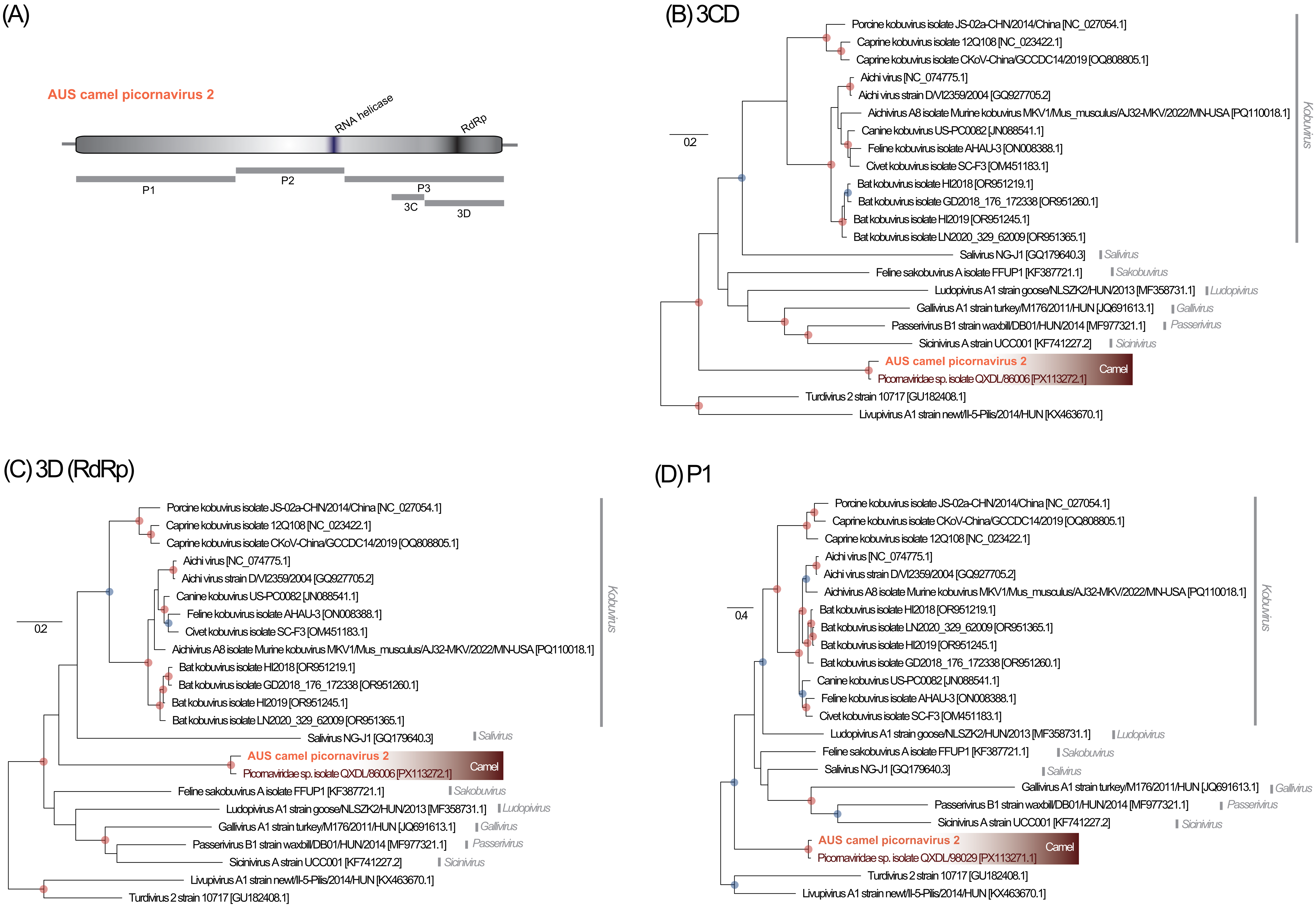

### Supplementary Figure S4

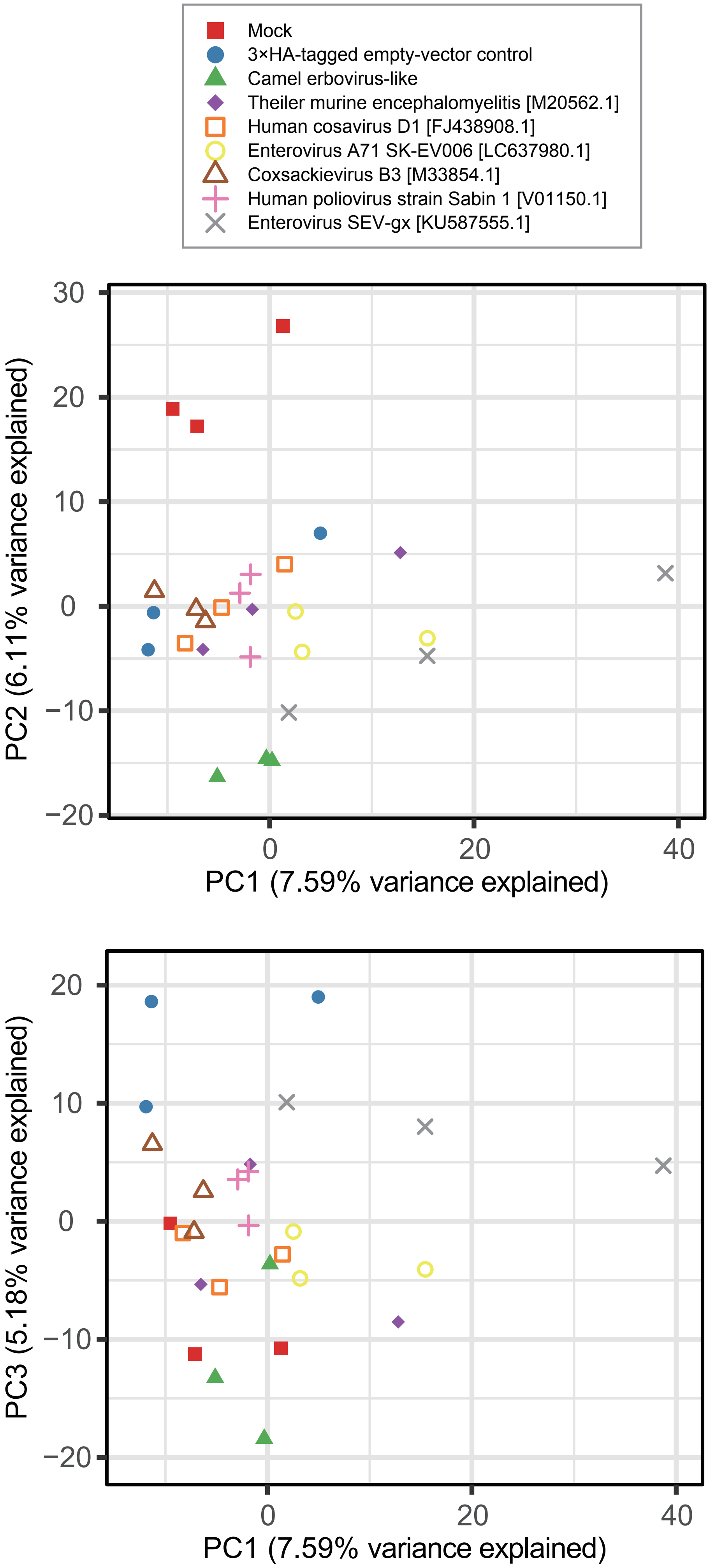
